# Magnetic tweezers meets AFM: ultra-stable protein dynamics across the force spectrum

**DOI:** 10.1101/2021.01.04.425265

**Authors:** Alvaro Alonso-Caballero, Rafael Tapia-Rojo, Carmen L. Badilla, Julio M. Fernandez

## Abstract

Proteins that operate under force—cell adhesion, mechanosensing—exhibit a wide range of mechanostabilities. Single-molecule magnetic tweezers has enabled the exploration of the dynamics under force of these proteins with subpiconewton resolution and unbeatable stability in the 0.1-120 pN range. However, proteins featuring a high mechanostability (>120 pN) have remained elusive with this technique and have been addressed with Atomic Force Microscopy (AFM), which can reach higher forces but displays less stability and resolution. Herein, we develop a magnetic tweezers approach that can apply AFM-like mechanical loads while maintaining its hallmark resolution and stability in a range of forces that spans from 1 to 500 pN. We demonstrate our approach by exploring the folding and unfolding dynamics of the highly mechanostable adhesive protein FimA from the Gram-positive pathogen *Actinomyces oris*. FimA unfolds at loads >300 pN, while its folding occurs at forces <15 pN, producing a large dissipation of energy that could be crucial for the shock absorption of mechanical challenges during host invasion. Our novel magnetic tweezers approach entails an all-in-one force spectroscopy technique for protein dynamics studies across a broad spectrum of physiologically-relevant forces and timescales.

## Introduction

Proteins that participate in processes such as mechanotransduction, cell adhesion, or muscle contraction are exposed and carry out their function under force^1,2^. Mechanical loads induce conformational changes on these proteins, triggering downstream events like the recruitment of molecular partners^3^, the increase of intermolecular bond lifetimes^4^, or the delivery of mechanical work^5^. These proteins exhibit different nanomechanical properties that determine the force magnitudes to which they respond, a feature which in turn is intimately linked to their function and is crucial for the processes they are involved with.

For more than 20 years, single-molecule force spectroscopy techniques have enabled the mechanical manipulation of proteins and provided us insightful views of mechanobiological processes at the nanoscale^6^. Atomic Force Microscopy (AFM), one of the first techniques developed and yet today among the most used ones, can apply well-calibrated forces to single proteins over a range that spans from 10 pN to 2000 pN^7,8^. AFM has been crucial to determine the nanomechanical properties of the Ig-like domains of the muscle protein titin^9–12^, the high mechanical stability of bacterial adhesion proteins^13–15^, or the extreme mechanostability of certain molecular interactions^16–18^, to name a few. No other single-molecule technique can reach the upper force ranges of AFM and explore ultra-stable folds and interactions. Unfortunately, although some successful examples exist^19,20^, its decreased resolution at low force ranges and instability have prevented the identification of protein folding events and slow-kinetic molecular events that occur at forces below 20 pN and in narrow ranges. By contrast, magnetic tweezers, since its first implementation for protein studies^21^, has demonstrated that its sub-pN resolution and week-long stability in the 0.1-120 pN range outcompete AFM for exploring protein dynamics at low forces and for extended periods^22–26^ (**Fig. 1**). Nevertheless, the inability to reach higher forces limits the extent of its application and prevents the exploration of highly mechanostable proteins and/or interactions. Therefore, there is no single-molecule force spectroscopy technique that provides high resolution and stability across the entire protein nanomechanical spectrum.

**Figure 1.**
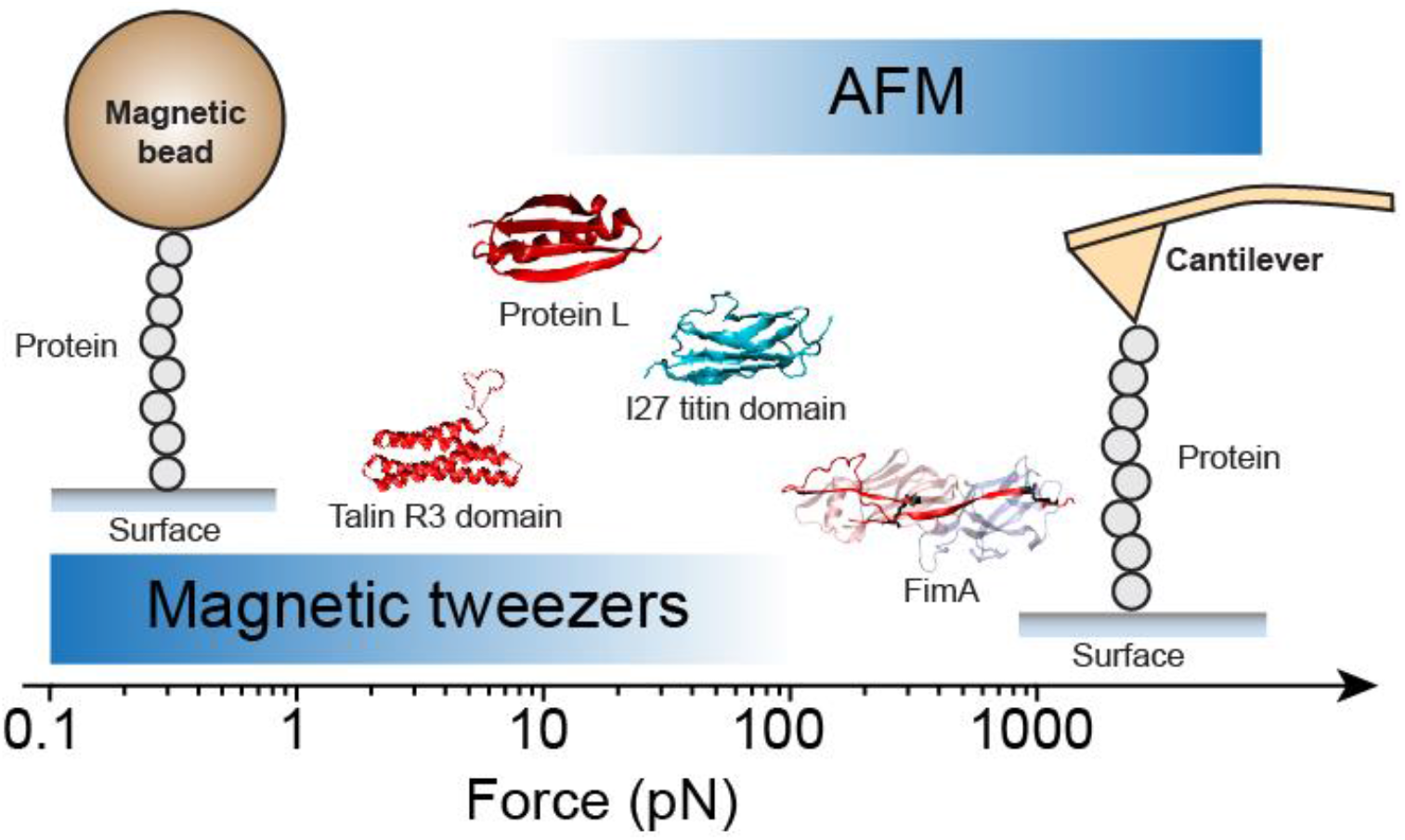
Magnetic tweezers *vs* AFM. Mechanostability varies greatly between proteins. The folding and unfolding dynamics of proteins like the talin R3 domain or protein L are better explored by magnetic tweezers (0.1-120 pN) due to its sub-pN resolution and stability, which permit to monitor their folding dynamics in narrow ranges (<1 pN) and over extended periods (>1 day). By contrast, mechanostable proteins like the I27 titin domain or the bacterial adhesion protein FimA are better approached with AFM (10-2000 pN), which can easily address their unfolding dynamics under high mechanical loads (PDB codes: R3, 2L7A; protein L, 1HZ6; I27, 1TIT; FimA, 3QDH).

Herein, we present a magnetic tweezers approach to achieve AFM-like forces while maintaining the idiosyncratic resolution and stability of magnetic tweezers. To reach higher forces, we implement, for the first time, the use and calibration of the superparamagnetic probes Dynabeads^®^ M-450 in two magnetic tweezers configurations: permanent magnets^23^ and magnetic tape head^25^. We achieve long-lasting experiments under high force by combining the HaloTag and split-protein techniques, which permit the end-to-end covalent tethering of proteins. To determine the force range accessible with the M-450 probes, we use the force-dependent extension changes and unfolding kinetics of model proteins to calibrate a force law for our two configurations. Our results indicate that, in the permanent magnet configuration, we can apply forces spanning from 1 to 510 pN, while in the magnetic tape head the force range spans from 0 to 236 pN. In both tweezers configurations, we can address the folding and unfolding dynamics of proteins of heterogeneous mechanostability while preserving subpiconewton, submillisecond, and nanometer resolutions. We demonstrate the capabilities of this upgrade by mapping the nanomechanics of the mechanostable type 2 pilus protein FimA, from the Gram-positive pathogen *Actinomyces oris*. FimA displays sharp asymmetric dynamics, with the unfolding occurring at loads >300 pN, while its folding proceeds at forces <15 pN. The broad force range separation between these two processes enables the dissipation of large amounts of energy, around 800 *k_B_T* per FimA subunit, which suggests that *A. oris* FimA type 2 pili could act as megaDalton-scale shock absorbers that protect bacterial adhesion from mechanical challenges. Despite the large increment in force that the M-450 superparamagnetic beads provide to magnetic tweezers, these probes can be manipulated with subpiconewton resolution to address the dynamics of proteins such as the mechanosensitive R3 domain from talin, whose folding equilibrium shifts in less than 1 pN. Our implementation of the M-450 probes unites the features of AFM and magnetic tweezers without sacrificing their strengths, and establishes an all-in-one force spectrometry technique for the study of most of the proteins and interactions across the force spectrum.

## Results

### Dynabeads^®^ M-450: double-covalent tethering and solid-phase assembly

Dynabeads^®^ M-450 are superparamagnetic microspheres of 4.5 μm in diameter commonly used for cell isolation. The forces that can be applied in single-molecule magnetic tweezers depend on the magnetic properties of the probe, and the strength and geometry of the magnets employed^27–29^. The M-450 beads show low bead-to-bead variation (1.2% variation in diameter) and, in comparison with the commonly used Dynabeads^®^ M-270, their volume is ~4 times larger and their magnetization moment almost twice^30^, which indicates that, under the same magnetic field, higher forces are accessible. In order to use the M-450 beads for magnetic tweezers, it is first required to obtain the specific and durable tethering of the molecule of interest, which constitutes a common challenge in the single-molecule force spectroscopy field^31–34^. Usually, mechanical loads exponentially decrease the lifetime of bonds^35^, and biomolecules exhibiting high mechanical stability are difficult to study because of the premature rupture of the linkage between the molecule of interest and the probe/surface employed in the single-molecule technique. We solved the specificity and the durability issues by using the HaloTag and split protein techniques, which allow the end-to-end covalent immobilization of the protein of interest between the glass surface of the experimental fluid chamber and the magnetic probe (**Fig. 2**).

**Figure 2.**
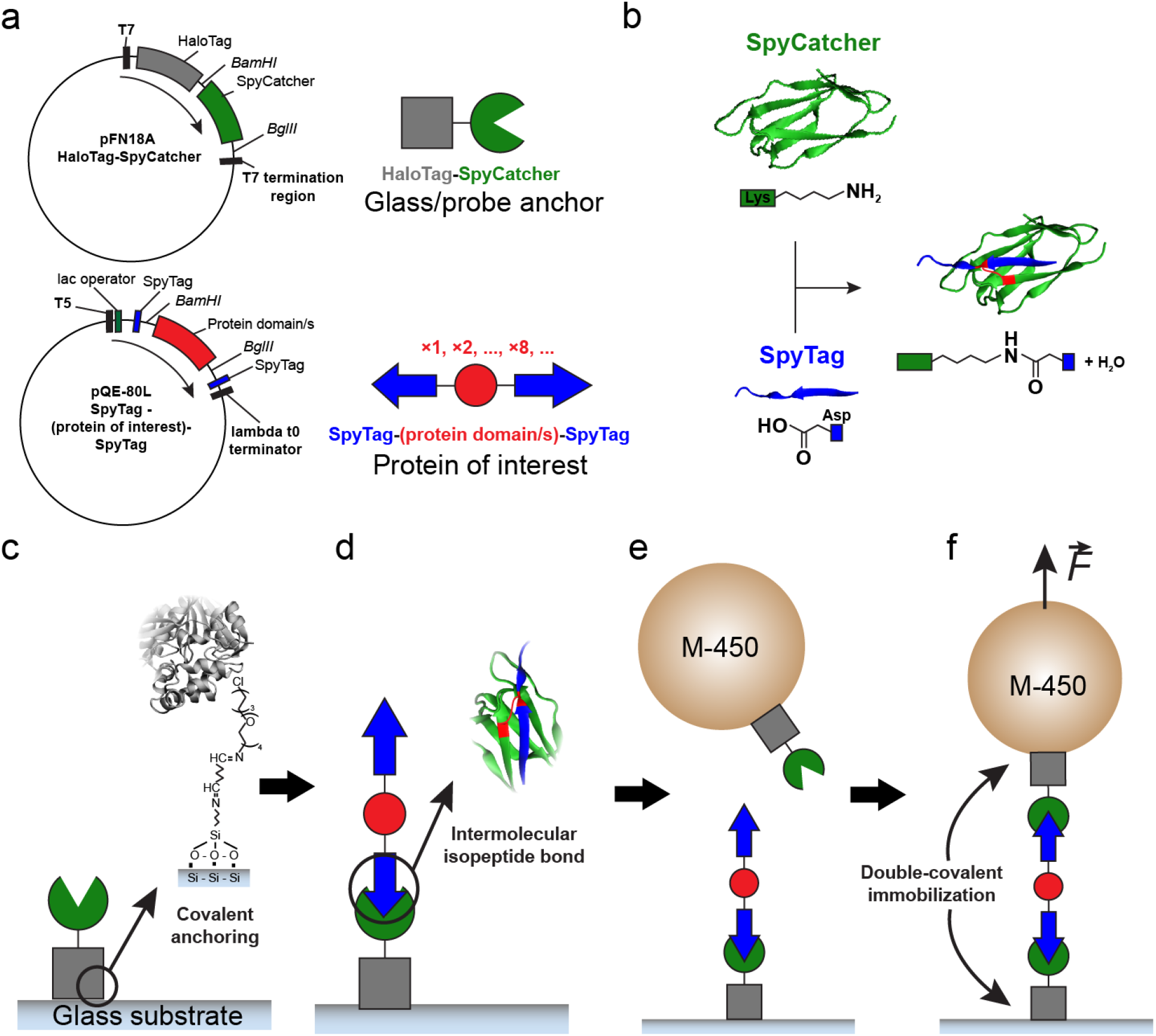
Double-covalent tethering and solid-phase assembly of proteins for single-molecule magnetic tweezers. **a)** Protein building blocks for double-covalent tethering and assembly for magnetic tweezers experiments. The glass and probe anchor module HaloTag-SpyCatcher is cloned in a modified pFN18A expression plasmid, while the protein of interest flanked by SpyTag peptides is cloned in a modified expression plasmid pQE-80L. In both plasmids, the gene sequences to be inserted are digested with the restriction enzymes BamHI-BglII, while the expression plasmid is only digested with BamHI (see **Supplementary Methods**) **b)** The split protein technique SpyCatcher/SpyTag. Each counterpart contains one of the residues that participate in the spontaenous formation of a covalent isopeptide bond. Specifically, SpyCatcher CnaB2 fold contains the reactive Lys31 and the catalytic Glu117 (not shown), while SpyTag peptide contains the reactive Asp117 residue. After reaction, Lys31 and Asp117 side chains form a covalent bond and a molecule of water is released. **c)** Protein assembly procedure for magnetic tweezers experiments. First, the anchor protein HaloTag-SpyCatcher (or HaloTag-SnoopCatcher) is covalently immobilized on the glass surface through the HaloTag. The glass substrate is functionalized to harbor the HaloTag ligand, which is bridged by glutaraldehyde to an APTES-modified glass slide. **d)** Second, the protein of interest (in red, it can be a single domain or a polyprotein), which is flanked by SpyTag (or SnoopTag) (in blue) and reacts with the SpyCatcher (or SnoopCatcher) forming an intermolecular isopeptide bond with the surface-anchored protein. **e)** M-450 probes are also functionalized with the anchor protein HaloTag-SpyCatcher. Functionalized beads are added to the fluid chamber where the SpyCatcher reacts with the free SpyTag from the surface protein assembly. **f)** Finally, the tethering and the assembly of the protein of interest is completed and its dynamics can be explored at high force.

The double-covalent anchoring method requires the engineering and cloning of two classes of protein modules; (1) the anchors, which bind covalently to the glass and M-450 surfaces, and the (2) proteins of interest, which constitute the molecule whose dynamics under force are to be studied (**Fig. 2a**). The anchor module is a HaloTag-based fusion protein that contains a SpyCatcher protein. This anchor is cloned and expressed from a modified pFN18A-HaloTag Flexi^®^ vector, and besides anchoring to the M-450 probe and glass surfaces through the HaloTag protein, it also covalently immobilizes the protein of interest. The protein of interest is cloned in a modified pQE-80L expression plasmid, and it can be either a single-domain or a polyprotein flanked by SpyTag peptides. When the SpyCatcher and the SpyTag partners are incubated together they bind by forming an intermolecular isopeptide bond, which reconstitutes the split-protein system (**Fig. 2b**). The isopeptide bond is formed between the side chains of a Lys residue—present in SpyCatcher— and an Asp residue—located in the SpyTag peptide. This spontaneous reaction proceeds efficiently, with high specificity, and under a wide range of conditions^36^. This split protein system has been extended to other molecular partners, such as SnoopCatcher/SnoopTag, of similar performance and specificity^37^, and which we also implement for some of our proteins (protein L octamer in **Fig. 3**, see **Supplementary Methods**). Hence, by engineering fused proteins containing the SpyCatcher and the SpyTag counterparts (or SnoopCatcher and SnoopTag), it is possible to covalently link with high specificity different polypeptide sequences and assemble molecular structures *à la carte*.

**Figure 3.**
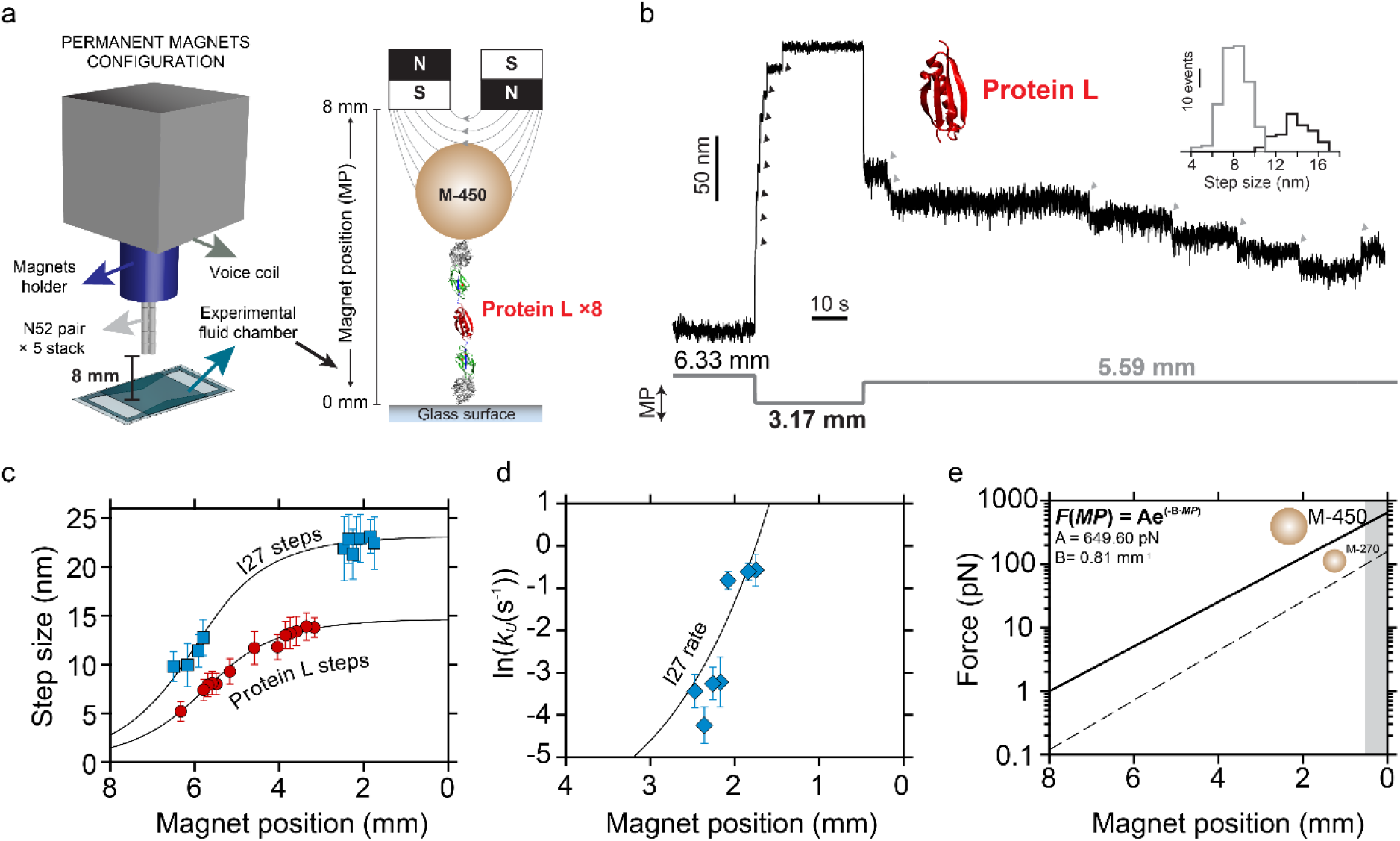
M-450 superparamagnetic bead force calibration for the permanent magnets configuration. **a)** In the permanent magnets configuration, the magnets are mounted on a voice coil device which moves them along a range of 8 mm, above the experimental fluid chamber (see **Supplementary Methods**). **b)** Force-clamp trajectory of an octamer of protein L. Along the experiment, the distance of the magnets is modified, which triggers protein unfolding (magnet position from 6.33 mm to 3.17 mm, black arrows), which results in step sizes of 13.8±1.4 nm (inset, black histogram, n=75), and folding and unfolding transitions (magnet position from 3.17 mm to 5.59 mm, grey arrows) which yield 8.1±1.2 nm step sizes (inset, grey histogram, n=191). **c)** Force-dependent extension changes (mean±SD) of folding/unfolding protein L (red circles) and I27 (blue squares) polyproteins as a function of the magnet position. **d)** Unfolding kinetics (mean±SEM) of I27 as a function of the magnets position. Extension changes and unfolding kinetics are fitted globally with the FJC (panel **c**, lines) and the Bell (panel **d**, line) models to obtain the parameters *A* and *B* (see panel **e**) that describe the magnet law that relates the magnets position (mm) with the force (pN) that can be applied with the M-450 beads. **e)** M-450 (solid line) and M-270 (dotted line) force range comparison along the 8 mm distance. Panel **c** data points for protein L step size: 6.33 mm, n=39; 5.79 mm, n=304; 5.69 mm, n=227; 5.59 mm, n=191; 5.49 mm, n=92; 5.17 mm, n=51; 4.59 mm, n=89; 4.04 mm, n=74; 3.85 mm, n=232; 3.74 mm, n=70; 3.59 mm, n=190; 3.36 mm, n=106; 3.17 mm, n=75. Data points for I27 step size: 6.5 mm, n=38; 6.17 mm, n= 55; 5.91 mm, n=35; 5.8 mm, n=48; 2.47 mm, n=35; 2.36 mm, n=23; 2.26 mm, n=15; 2.17 mm, n=45; 2.08 mm, n=55; 1.84 mm, n=23; 1.75 mm, n=20. Panel **d** data points for Data points for I27 unfolding kinetics: 2.47 mm, n=27; 2.36 mm, n=20; 2.26 mm, n=14; 2.17 mm, n=45; 2.08 mm, n=55; 1.84 mm, n=22; 1.75 mm, n=17.

To assemble our molecular structures for magnetic tweezers experiments, we utilize a solid-phase approach that involves the sequential addition of protein blocks. Starting from the glass surface of the fluid chamber, we first immobilize the protein anchor HaloTag-SpyCatcher. As previously described^23,33,38^, we functionalize the glass surface with the HaloTag ligand, which is recognized and covalently bound by the HaloTag protein (**Fig. 2c**). After the immobilization of the anchor, we incubate the second module of the assembly, the protein of interest, on the glass surface. The SpyTags located on the N or C-termini of the protein of interest can bind to the SpyCatcher from the surface-immobilized anchors, covalently anchoring the protein (**Fig. 2d**). By incubating the glass surface with very low concentrations of the first module—protein anchor— we minimize the occurrence of surface bridges, where both SpyTag peptides of the same protein of interest end up connected to two glass-side protein anchors. In the final step, we close the assembly with a third module of HaloTag-SpyCatcher, which has been previously immobilized on the surface of the M-450 bead. The tosylactivated surface of the M-450 allows the binding of primary amine and thiol groups, which facilitates the fast and easy covalent anchoring of the protein anchors. We circulate over the glass surface the functionalized M-450 beads, whose HaloTag-SpyCatcher anchors will recognize and bind to the SpyTag present on the free end of the protein of interest (**Fig. 2e**). This last reaction closes the assembly and generates a double-covalent tethered molecule ready to be tested in magnetic tweezers (**Fig. 2f**). With this solid-phase approach, the sequential addition of modules permits the assembly of protein tethers for their study under high mechanical loads for days-long periods. The formation of the intermolecular covalent bond prevents the mechanical extension of the reconstituted SpyCatcher/SpyTag, which provides an inextensible handle during the force spectroscopy experiment

### Dynabeads^®^ M-450 force calibration in the permanent magnets configuration

In our permanent magnets configuration (**Fig. 3a**), the fluid chamber containing the tethered protein of interest is mounted over the objective of an inverted microscope. Above the fluid chamber, the position of a pair of N52 neodymium magnets (D33, K&J Magnetics) is controlled with a voice-coil actuator (LFA-2010, Equipment Solutions) operated under feedback. The voice-coil moves the magnets vertically, approaching or moving them away from the sample along a distance of 8 mm. To elucidate the forces that we can apply with the M-450 beads, we use an empirical magnet law that permits us to establish a relationship between the magnets position (*MP*) and the force being applied to the protein under study (*F*)^23^:

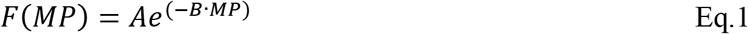

where *MP* is the magnet position in mm, and *A* and *B* are the fitting parameters of the exponential magnet law, in pN and mm^−1^, respectively. In order to obtain a force calibration curve, it is required to measure an observable that changes with the mechanical load. As we previously demonstrated^23,25^, the force-dependent extension changes of an unfolding or folding protein (step size) accurately follow standard polymer models such as the freely-jointed chain (FJC) model^39^, hence, constituting a very sensitive gauge. However, this force scaling is highly non-linear, resulting in very small extension changes at forces above 50 pN that can lead to an inaccurate calibration at high forces. By contrast, the unfolding rate of a protein increases exponentially with the mechanical load, providing a very sensitive observable in the high force range that can be described by the Bell model^35^. Therefore, to obtain an accurate magnet law over the complete force range, we opt for a combined approach by simultaneously utilizing both observables to calibrate the force-magnet position dependency (Eq. 1): first, to asses the low force regime, we employ the step sizes of two model proteins, the bacterial superantigen protein L and the I27 titin domain, whose force dependencies have been thoroughly measured both with magnetic tweezers and AFM^9,23,33,40,41^; second, to account for the high force regime, we use the unfolding kinetics of the mechanostable I27, previously measured with AFM and described with the Bell model^42^. Thanks to this combined approach, which takes advantage of the opposite force-sensitivity of two different observables at low and high forces, we can do a joint fit to Eq. 1 to obtain an accurate estimation of the *A* and *B* parameters and, hence, a precise calibration across the full force spectrum.

**Figure 3b** shows a trajectory of the dynamics under force of an octamer of protein L, which embodies the first part of the calibration. Initially, at *MP*=6.33 mm all of the domains remain in the folded state. After the approximation to *MP*=3.17 mm, the unfolding of the eight protein L domains is rapidly observed as 13.8±1.4 nm (mean±SD) step wise extensions. Once the magnets are moved away to *MP*=5.59 mm, the protein elastically recoils and the domains start to undergo transitions between the folded and unfolded states, yielding step sizes of 8.1±1.2 nm. In this manner, we obtain the force-dependent step sizes of the folding and unfolding transitions of protein L and I27 (**Supplementary Fig. 1**) as a function of the magnet position (**Fig. 3c**). These folding/unfolding events yield step sizes (*x*) whose values scale with the force applied (*F(MP)*) and can be described by the freely jointed chain polymer model^39^ (FJC):

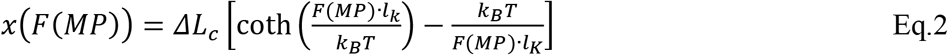

Where *ΔL_C_* is the contour length change in nm, *lK* is the Kuhn length in nm, and *k_B_T* is the thermal energy as 4.11 pN∙nm. The measurement of the force-dependent folding/unfolding step sizes of these two proteins proves that, despite their larger size, the use of the M-450 beads does not entail a loss of resolution since it is possible to resolve the step size variation with nanometer resolution.

For the calibration of the high force regime, we measure the unfolding dwell-times^43^ of I27 (**Supplementary Fig. 1**) as a function of the magnet position (**Fig. 3d**), and we use the Bell model for bond lifetimes under force to describe this dependency^35^:

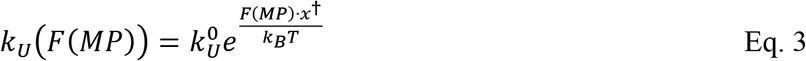

Where *k^0^_U_* is the extrapolated unfolding rate in the absence of force in s^−1^, *x^†^* is the distance to the transition state in nm, *k_B_T* is thermal energy as 4.11 pN∙nm. Finally, we obtain the magnet law for the M-450 beads by fitting simultaneously (see **Supplementary Table 1**) our three observables: the step sizes of protein L and I27—fitting Eq. 2—, and the unfolding kinetics of I27—fitting Eq. 3. The step sizes of both proteins and the unfolding kinetics of I27 are accurately described by this approach (solid lines in **Fig. 3c** and **d** graphs), and it allows us to determine the values of the *A* (649.60±119.37 pN) and *B* (0.81±0.09 mm^−1^) parameters of the magnet law from Eq. 1. This magnet law, represented in **Fig. 3d**, indicates that the forces we can apply to proteins span from 1 pN (8 mm) to 649 pN (0 mm). However, the maximum force of application is ~510 pN, since the thickness of the experimental fluid chamber prevents the approximation of the magnets to positions <0.3 mm (shaded area in **Fig. 3d**). The increase in the upper force range obtained with the M-450 beads is more than 4 times larger than with the M-270, which enables the unfolding of highly mechanically stable proteins. This force calibration indicates that, for the first time, forces previously only attainable with AFM, can now be used in magnetic tweezers, while maintaining its resolution and stability.

### Dynabeads^®^ M-450 force calibration in the magnetic tape head configuration

Magnetic fields can be generated either by a pair of permanent magnets or by an electromagnet. In the latter, the intensity of the magnetic field—and hence the force—is modulated by the current that is passed through a coil wrapped around a magnetic core. In this second configuration of magnetic tweezers, we use a magnetic tape head (902836, Brush Industries) to apply forces to the molecules under study. This configuration presents some advantages in comparison to the permanent magnet one, like the ability to change the force faster (100 ms *vs* 40 μs), the increased stability, and the higher signal-to-noise ratio. Because of these advantages, the use of this magnetic tweezers configuration is best suited for identifying short-lived molecular conformations^25^, ligand-binding events^44^, or exploring protein dynamics under complex force signals^45^. As we previously showed ^25^, in this configuration the force (*F(I, z)*) depends on the current (*I*) and the distance between the head and the bead (*z*). In our card-reader design (**Fig. 4a**), the magnetic tape head is held fixed at *z*=300 μm (see **Supplementary Methods**), therefore the force only depends on the current (*F(I)*):

**Figure 4.**
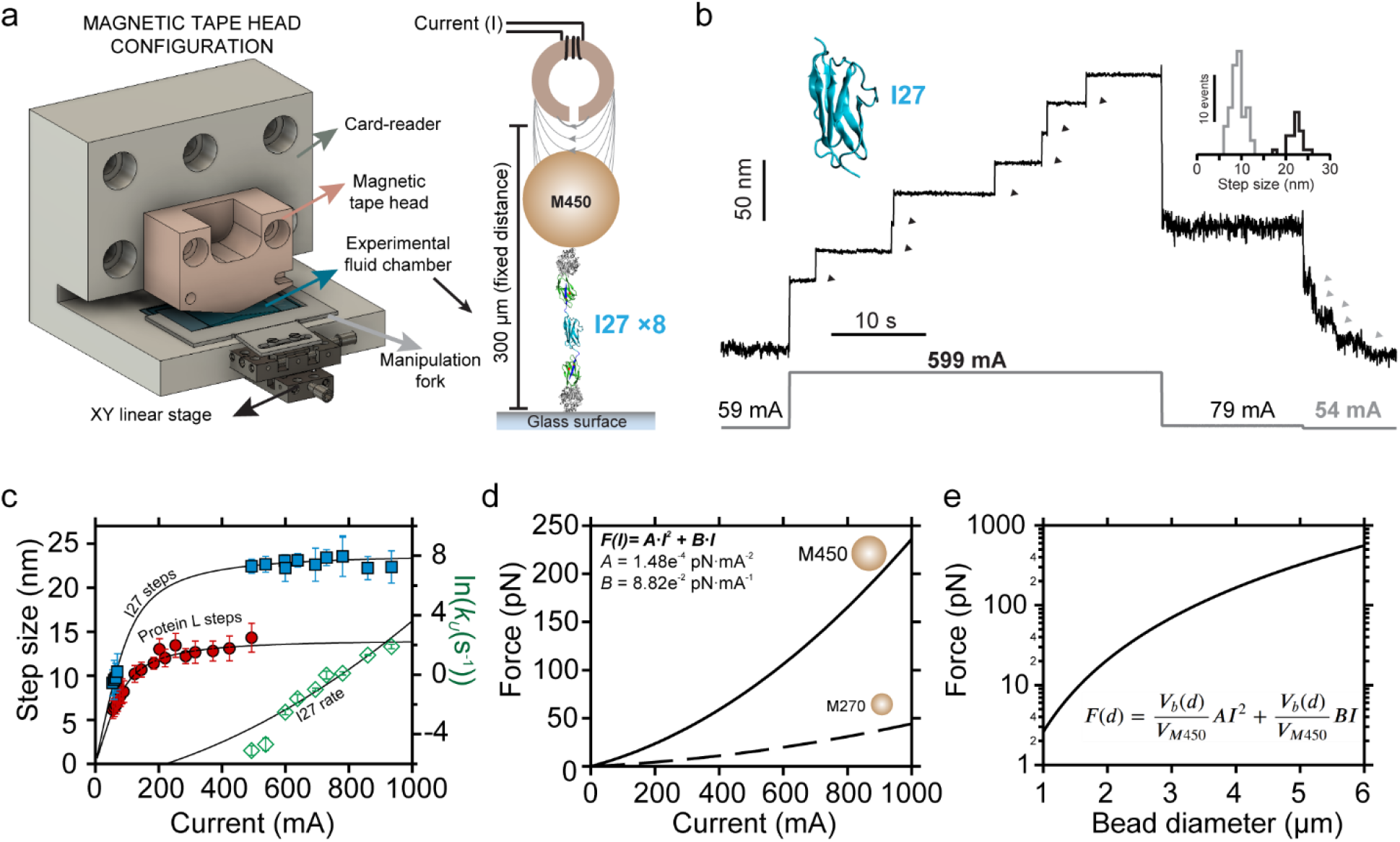
M450 superparamagnetic bead force calibration for the magnetic tape head configuration. **a)** In the card-reader configuration, the magnetic tape head is held fixed 300 μm above the sample, and the force is modulated by the current (mA) passed through the head. **b)** Force-clamp trajectory of an octamer of I27. Along the experiment, the current is modified, which triggers protein unfolding (current value from 59 mA to 599 mA, black arrows), which results in step sizes of 22.2±1.5 nm (mean±SD, inset, black histogram, n=25), and folding transitions (current value from 79 mA to 54 mA, grey arrows) which yield 9.2±1.5 nm step sizes (inset, grey histogram, n=78). **c)** Force-dependent extension changes (mean±SD) of folding and unfolding protein L (red circles) and I27 (blue squares) polyproteins, and the unfolding kinetics (mean±SEM) of I27 (green diamonds), as a function of the current (mA). For the polymer extensions we use the FJC model for polymer elasticity, and for the unfolding kinetics we assume the Bell model for bond lifetimes. Both observables—polymer extension and unfolding kinetics—are fitted simultaneously to obtain the parameters *A* and *B* (see panel **d**) that describe the current law that relates the current value (mA) with the force (pN) that can be applied with the M450 beads. **d)** M-450 (solid line) and M-270 (dotted line) force range comparison from 0 to 1000 mA. **e)** Force magnitude as a function of the bead diameter. Assuming the properties of the M-450 and normalizing by their volume, beads of increasing diameter up to 6 μm can give access to forces above 550 pN. Panel **c** data points for protein L step size: 55 mA, n=51; 60 mA, n=17; 65 mA, n=63; 70 mA, n=122; 75 mA, n=76; 79 mA, n=84; 84 mA, n=30; 89 mA, n=92; 124 mA, n=33; 145 mA, n=38; 184 mA, n=41; 202 mA, n=59; 220 mA, n=137; 253 mA, n=72; 285 mA, n=183; 315 mA, n=107; 371 mA, n=169; 423 mA, n=137; 493 mA, n=20. Data points for I27 step size: 54 mA, n=78; 60 mA, n=9; 65 mA, n=14; 70 mA, n=6; 493 mA, n=11; 599 mA, n=39; 638 mA, n=15; 694 mA, n=93; 729 mA, n=6; 780 mA, n=95; 859 mA, n=28; 934 mA, n=32. Data points for I27 unfolding kinetics: 493 mA, n=11; 599 mA, n=39; 638 mA, n=15; 694 mA, n=89; 729 mA, n=6;780 mA, n=95; 859 mA, n=28; 934 mA, n=32.

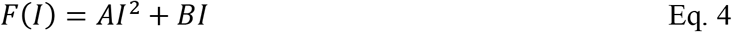

Where *A* (pN∙mA^−2^) and *B* (pN∙mA^−1^) are the fitting parameters.

Following the same procedure as with the permanent magnets, we calibrate the current law by measuring the force-dependent folding/unfolding step sizes of protein L and I27, and the unfolding kinetics of I27, as a function of the current. **Figure 4b** shows a trajectory of an octamer of the I27 titin domain. At 59 mA, seven out of eight domains of I27 remain in the folded state. After increasing the current to 599 mA, the unfolding extensions of the seven I27 domains appear as 22.2±1.5 nm length steps. Changing the current value to 79 mA permits the elastic contraction of the unfolded polyprotein, but not the folding of the I27 domains, which finally are detected as fast contraction steps (9.2±1.5 nm) after decreasing the current to 54 mA. Hence, we measure the force-dependent extension changes of folding and unfolding I27 and protein L (**Supplementary Fig. 2**) and the force-dependent unfolding kinetics of I27. We use equivalent expressions to the ones in Eq. 2 and Eq. 3, but considering the force as a function of the current value:

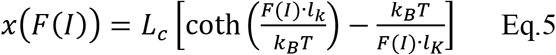

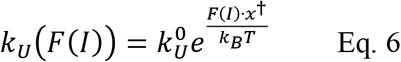

We fit simultaneously (**Fig. 4c**) these three observables and obtain the values of *A* = (1.48 ± 0.12)×10^−4^ pN∙mA^−2^ and *B* = (8.83 ± 0.98)×10^−2^ pN∙mA^−1^ for the current law of Eq. 4 (**Supplementary Table 2**). **Fig. 4d** compares the current law for the M-450 and the M-270 beads. With the M-450, the range of forces spans from 0 (0 mA) to 236 pN (1000 mA), which entails an increment of more than five times the maximum force attainable with the M-270 beads (0-40 pN). **Supplementary Fig. 3** shows an analytical prediction of the force-current law for the superparamagnetic beads MyOne, M-270 (experimentally determined^25^), and M-450 (theoretical and experimental), based on their properties and the properties of our magnetic tape head^25,30^. The experimentally determined relationship between the current and the force, based on our measurements of protein extension changes and unfolding kinetics, matches the theoretical one for the M-450 beads, which supports the strength of our calibration method. Despite the large increase in force, in the magnetic tape head configuration we can apply half the force magnitudes that we can use in the permanent magnets. In order to reach higher forces with this same magnetic tape head, the sample should be brought closer to the head gap (less than 300 μm) or implement larger beads or beads exhibiting higher magnetization properties. Considering the bead size, **Fig. 4e** shows a prediction of the force range accessible with the magnetic tape head configuration as a function of the diameter of a putative superparamagnetic probe that exhibits the same properties as the M-450. Under the magnetic field induced by a current of 1000 mA, beads of 6 μm in diameter would permit to apply more than 550 pN. As new superparamagnetic beads become commercially available, our magnetic tape head configuration and our calibration method offer the perfect theoretical and experimental framework for future force calibrations of both smaller and larger probes.

### FimA pilus protein dynamics in magnetic tweezers

The implementation of the M-450 superparamagnetic beads provides a significant leap in the force range that can be used in our two magnetic tweezers configurations. We challenged the covalent anchoring strategy and the newly-achieved upper force ranges of magnetic tweezers by exploring the dynamics of the highly mechanostable protein FimA, from the type 2 pili of the dental plaque Gram-positive pathogen *Actinomyces oris*. Previously, we explored with AFM the dynamics of FimA, which showed rupture forces ~700 pN when stretched under constant speed conditions (400 nm∙s^−1^)^14^. The high mechanostability of FimA is partly due to the presence of two strategically-located intramolecular isopeptide bonds, which prevent the mechanical extension of most of the FimA structure^46^. These bonds enclose a sequence of 44 residues that define the isopeptide-delimited loop (IDL) motif, which is the only part of FimA that can extend under force.

To explore the folding and unfolding dynamics of FimA, we use the permanent magnets configuration, which can apply forces in the 1-510 pN range. We designed a polyprotein made of four tandem repeats of the CnaA-N2/CnaB-N3 domains of FimA, flanked by SpyTag sequences. This single-polypeptide made of four FimA subunits evokes the architecture of a pilus and maintains the nanomechanical properties that derive from its assembly (see **Supplementary Fig. 4**). We covalently immobilized end-to-end the FimA polyprotein by anchoring its N and C-terminal SpyTags to HaloTag-SpyCatcher proteins bound to the glass and the M-450 bead surfaces (**Fig. 4a**).

The trajectory represented in **Fig. 5b** shows the folding and unfolding dynamics of the FimA polyprotein. Forces >300 pN trigger FimA IDL unfoldings, which are detected as four 12.2 ± 1.3 nm discrete steps. Subsequently, we quench the force to 12.2 pN, where we can detect, for the first time, discrete 8.3 ± 2.3 nm folding steps corresponding with the contraction of the four previously unfolded FimA IDLs. In this manner, triggering the unfolding and folding at different forces of FimA IDL we characterize the nanomechanical properties of this protein, such as its force-dependent folding/unfolding extension changes (**Fig. 5c**) and folding probability (**Fig. 5d**). IDL folding probability plot shows that in a 3 pN range the protein switches from totally unfolded (~14 pN) to completely folded (~11 pN), which resembles the behavior observed in Ig-like domains with engineered disulfide bonds^47^. We characterize the folding/unfolding kinetics of FimA IDL in the 6-420 pN range (**Fig. 5e**). We find that, in the absence of force, FimA IDL folding (using the rate equation used in references^47–49^) proceeds at rates of 4.44 ×10^4^ s^−1^, while its unfolding (using the rate equation based from the Bell model^35^) is predicted to occur at 2.89 ×10^−8^ s^−1^. Unlike unfolding (**Fig. 5e** square symbols), the folding rate of FimA (**Fig. 5e** arrow symbols) shows a strong force-dependency. This pronounced asymmetry (blue and red fits in **Fig. 5e**) highlights the different sensitivity to the mechanical load of these opposed transitions.

**Figure 5.**
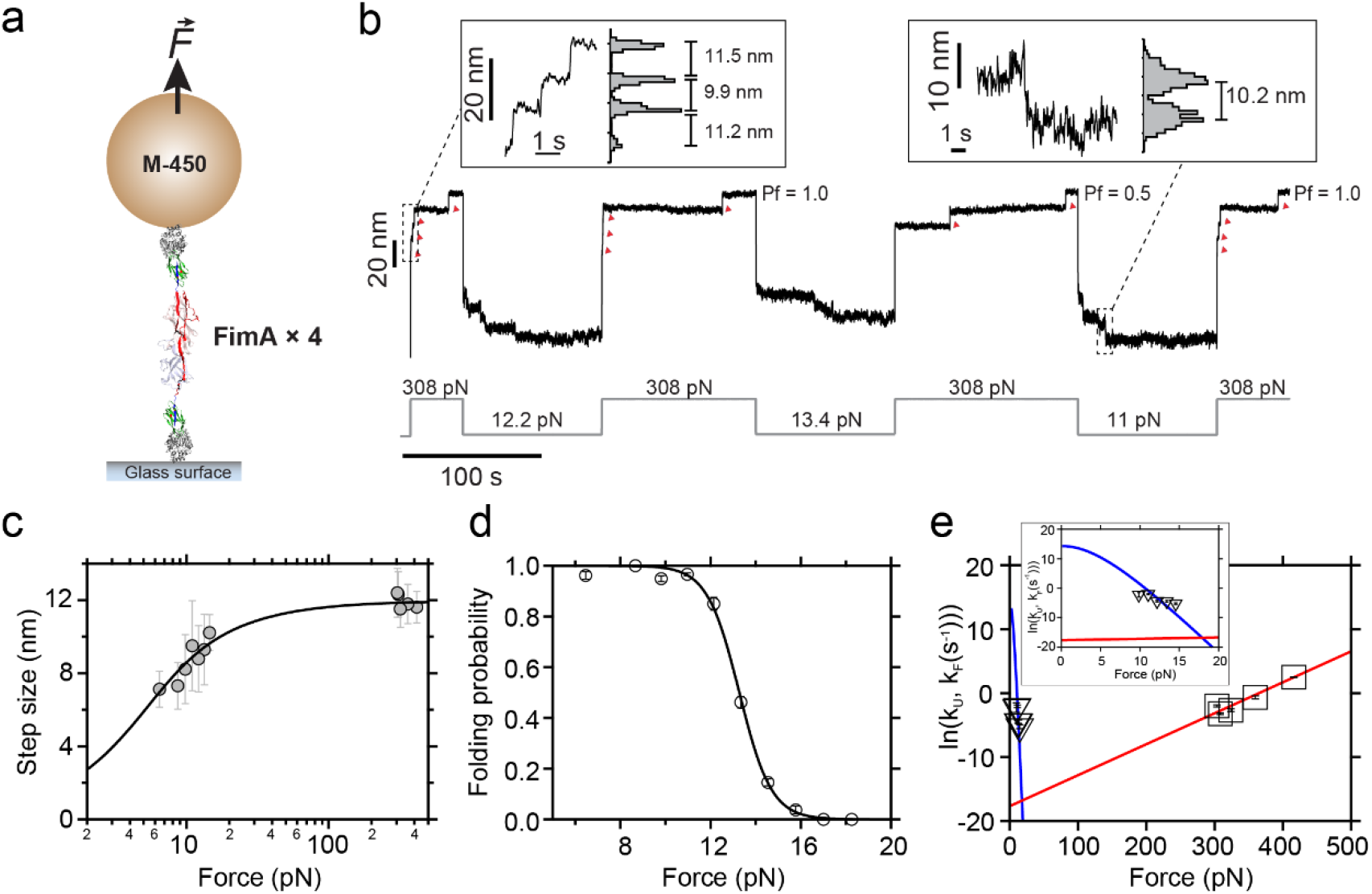
Dynamics under force of *A. oris* FimA pilin protein. **a)** In the permanent magnets configuration, we use a polyprotein made of four copies of FimA flanked by SpyTags sequences and covalently anchored to the glass substrate and the M-450 bead by HaloTag-SpyCatchers. **b)** Force-clamp trajectory of the FimA polyprotein. Forces >300 pN unfold FimA IDLs in less than 200 s, yielding step sizes ~12 nm. Onthe contrary, FimA IDL folding only occurs when the force is quenched to values <15 pN. **c)** IDL extension (mean±SD) as a function of the force. The line represents a fit to the data of the freely jointed chain model for polymer elasticity *ΔL_c_*= 11.97 nm, *l_k_*=1.44 nm (6.5 pN, n=32; 8.7 pN, n=17; 9.8 pN, n=28; 11.0 pN, n=58; 12.1 pN, n=24; 13.3 pN, n=19; 14.5 pN, n=6; 303 pN, n=154; 308 pN, n=170; 318 pN, n=79; 359 pN, n=34; 416 pN, n=42). **d**) FimA IDL folding probability (mean±SD). The line is a sigmoidal fit of the data points (6.5 pN, n=8; 8.7 pN, n=8; 9.8 pN, n=11; 11.0 pN, n=17; 12.1 pN, n=11; 13.3 pN, n=15; 14.5 pN, n=16; 15.8 pN, n=8; 17.0 pN, n=4; 18.3 pN, n=1). **e)** FimA folding and unfolding (mean±SEM) kinetics (9.8 pN, n=16; 11.0 pN, n=42; 12.1 pN, n=16; 13.3 pN, n=7; 14.5 pN, n=5; 303 pN, n=154; 308 pN, n=170; 318 pN, n=79; 359 pN, n=34; 416 pN, n=42;). The blue line is a fit to the IDL refolding kinetics (*ΔL_c_*= 11.97 nm, *l_k_*= 1.44 nm, ln*k_F_^0^*= 14.32) using the method described in references 47-49, and the red line is a Bell model fit (ln*k_U_^0^*= −17.67, *x_U_^†^*= 0.21 nm) of the IDL unfolding rate data.

Our measurements reveal the large gap between the forces at which FimA folding and unfolding occur. From the biological point of view, this hysteresis could act as a dissipation mechanism of the force-related stresses that bacteria endure during host colonization. Being an oral pathogen, an organism like *A. oris* could benefit from this biomechanical trait to withstand the forces exerted by mastication or brushing. We quantify the dissipative capacities of this pilus protein by applying a force-ramp (±17 pN∙s^−1^) stretching-relaxation protocol (**Supplementary Fig. 5a**) that triggers unfolding and folding on FimA polyproteins. The heat dissipated during this force-ramp protocols is the area enclosed by the stretching—high force—and relaxation—low force— curves (**Supplementary Fig. 5b**). During these measurements, we observe the unfolding at high force of the FimA domains and the subsequent contraction of this extended polymer as the load is decreased. The FimA domains only fold when the force is very low, and this situation generates a large hysteresis between both curves. Our results indicate that, at a load rate of 17 pN∙s^−1^, each FimA can dissipate ~800 *k_B_T* and this contribution adds up linearly with the number of unfolding FimA subunits in the polyprotein (**Supplementary Fig. 5b** and **c**, **Supplementary Methods**). If we consider that a single type 2 pilus is made of 100-150 FimA subunits, the energy dissipated by one of these fibers could oscillate between 7.0-12.5×10^4^ *k_B_T*, converting these structures in megaDalton-scale shock absorbers.

### From hundreds of piconewtons to subpiconewton ranges: the mechanosensitive talin R3 domain

The mechanical unfolding of FimA proves that magnetic tweezers now can operate at AFM-like force ranges and gives access to study highly stable molecules. Nevertheless, the bulkier dimensions of the M-450 beads, in comparison with the magnetic probes MyOne or M-270, could entail a potential loss in resolution and sensitivity that hinder the precision with which force can be manipulated in magnetic tweezers. To address this question, we measure the folding dynamics of the mechanosensitive R3 domain of the focal adhesion protein talin in our magnetic tape head configuration. Under force, the mutant R3^IVVI^ protein exhibits bistability in the ~7.5-9.5 pN range, experiencing an 80% shift in its folding equilibrium within less than 1 pN. We synthesize the protein HaloTag-R3^IVVI^-(Spy0128)_2_-SpyTag which directly binds to the glass surface through its HaloTag and which contains two copies of the pilin protein Spy0128 from *Streptococcus pyogenes*, which serve as an inextensible molecular handle^13^ (**Fig. 6a**). **Fig. 6b** shows trajectories of the R3^IVVI^ protein undergoing folding and unfolding at different force values 0.3-0.8 pN apart, from which we obtain the residency time at each state. As it can be seen in **Fig. 6c**, we can resolve the folding dynamics of the talin R3^IVVI^ domain in the ~8.0-10.0 pN range with the same resolution as previously reported with the M-270 beads^22,25,44,45^, demonstrating that the M-450 beads are a robust magnetic probe that permits to address molecular processes with high resolution under a wide range of physiological timescales and forces.

**Figure 6.**
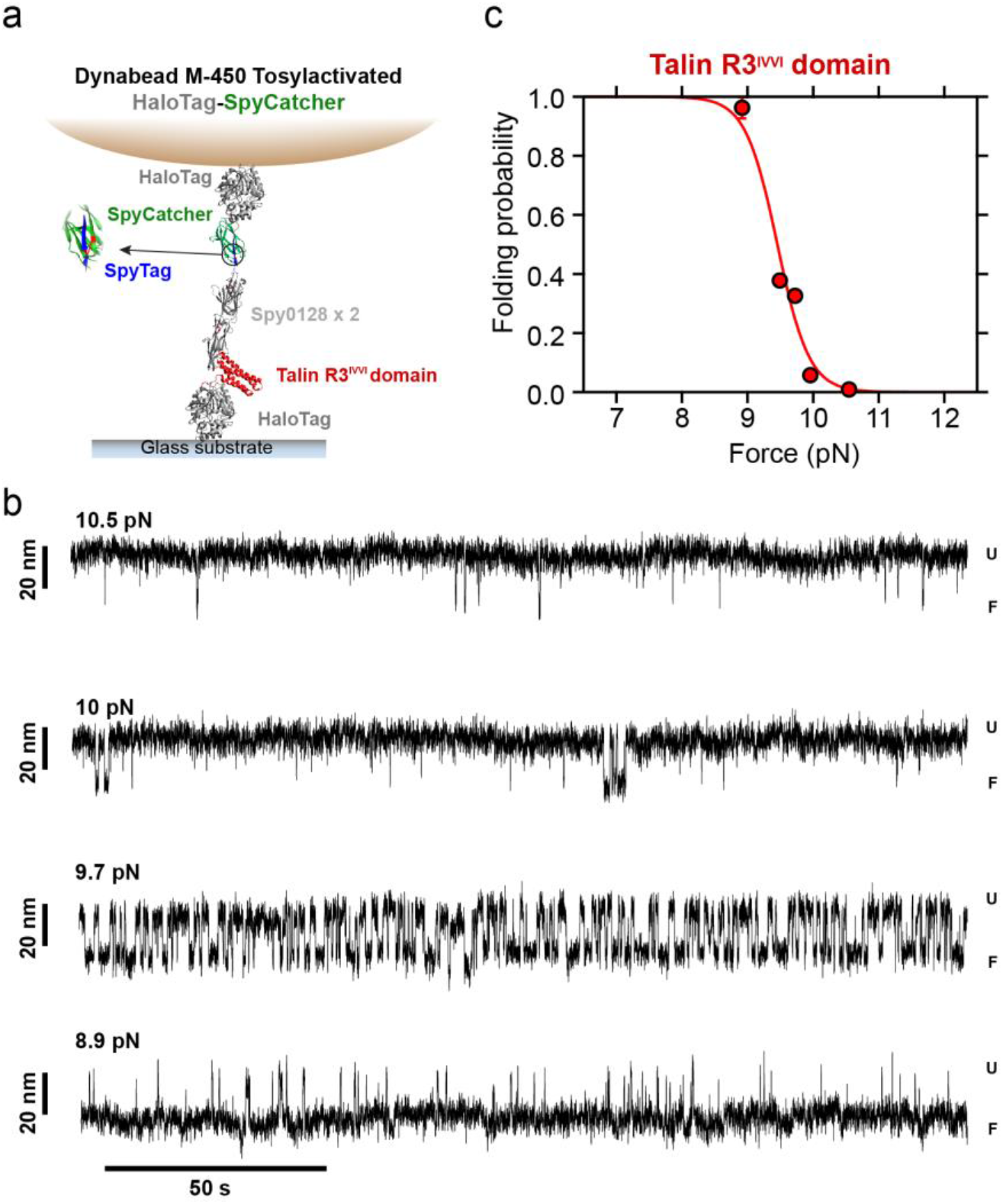
Talin R3^IVVI^ domain folding dynamics in the magnetic tape head configuration. **a)** Scheme of the covalent thetering of the single-domain protein of interest talin R3^IVVI^ domain. Unlike the other proteins used in this research, this chimeric protein sequence from N-to-C terminus is HaloTag-R3^IVVI^-(Spy0128)_2_-SpyTag. Spy0128 is the pilin protein of *Streptococcus pyogenes* and the two copies in our construct act as inextensible linkers. The M-450 bead is functionalized with HaloTag-SpyCatcher which recognizes and covalently binds to the SpyTag peptide present in the C-terminus of the protein of interest. **b)** Folding equilibrium trajectories of R3^IVVI^ domain registered in a force range window of 1.5 pN. This protein exhibits a bistable behavior at ~9-10.5 pN where the domain undergoes transitions between the unfolded (U) and folded (F) states. **c)** Folding probability of talin R3^IVVI^ domain. Each point represents the occupancy time ratio and the SD. Solid line is a sigmoidal fit to the data.

## Discussion

The effect of force on proteins governs a myriad of key biological processes. Mechanical perturbations not only affect proteins with mechanical roles, but also to all proteins along their lifetime, such as when they are synthesized, translocated, or degraded^1^. Single-molecule force spectroscopy techniques have enabled to manipulate single molecules and have revolutionized our understanding of protein dynamics. AFM, optical tweezers, or magnetic tweezers have permitted to delve into this topic and have provided a wealth of information^6^. From these studies, we know now that while some proteins undergo dramatic conformational changes within less than 1 pN^22,24,25^, some other proteins or protein-protein interactions unfold or break when exposed to hundreds of pN^17^, and only fold or reform at very low mechanical loads. This wide nanomechanical gap has challenged the usability of each of the aforementioned techniques. In AFM, this issue has been addressed with the development of probes with different stiffness, mainly oriented to enhance the resolution at low force by decreasing the spring constant of the cantilevers. However, this approach again entails a compromise and a limitation; although the resolution at low force can be increased, these soft cantilevers perform poorly and are driven out of range when used at high forces. Therefore, exploring the dynamics of a protein across its entire range of physiological forces has been prevented by the lack of a technique that grants access to the whole force spectrum. Here, we have demonstrated a gap-joining approach that pushes magnetic tweezers to the AFM force range playground, without compromising any of the technical advantages offered by this technique. With the same probe, we have sub-pN resolution and full control of the force across a range that spans from 0 to 520 pN, which permits to address the folding and unfolding dynamics of the majority of proteins.

Although the force range of magnetic tweezers now overlaps with that of the AFM and same molecular properties can be measured (**Supplementary Fig. 6**), both techniques exhibit differences in their mode of operation that makes them complementary and more or less suitable depending on the task. For instance, in AFM it is possible to sample hundreds of molecules per day, which enables to map with precision the dynamics of proteins with unbeatable efficiency. This high-throughput nature permits to easily detect conformational heterogeinities^50^ or probe the chemical diversity of molecules, such as the effect of small chemical modifications on the side chains of residues directly involved in the protein’s mechanical stiffness^51–53^, or to probe the effect of small molecules or peptides on the mechanical stability of proteins^54,55^. Magnetic tweezers measurements, by contrast, usually involve the testing of a lower number of molecular entities, although efforts towards parallelization have been successful^24,56^. Unlike in AFM, each of these molecules can be explored for periods that span from minutes to days^23,24^. The hallmark stability of magnetic tweezers grants access to rare events that manifest over slow timescales (>hours), such as irreversible chemical modifications^57^, and enables to directly map the nanomechanical properties of proteins evolving with very slow folding kinetics^23,24,40,49^. Moreover, magnetic tweezers offers the possibility of manipulating the force and the measurement conditions in real-time, which places this technique as the ideal platform for monitoring mechanochemical reactions or the effect of protein-protein interactions by iteratively changing the buffer composition^26,44,47^. Notably, while the limited time resolution was an initial shortcoming in magnetic tweezers, recent advances have expanded the technique to the kHz range^58^. This, combined with the high stability of magnetic tweezers, brings new challenges in data manipulation and analysis, as a single protein recording can comprise over 9×10^8^ data points (1 week at 1.5 kHz), bridging the single molecule with the Big Data field. As new technical improvements allow to increase the resolution and parallelization of the measurements, the development of new computational methods will be required for the handling of these massive amounts of data.

While both techniques exhibit features that makes them more or less adequate depending on the question, our force calibration with the M-450 beads proves that highly stable proteins, previously unreachable with magnetic tweezers, are now accessible with this technique. The protein FimA, from the adhesive pili of the Gram-positive pathogen *A. oris*, one of the causative agents of dental plaque^59^, is the archetype of high mechanical stability. Our previous AFM work, addressed the unfolding dynamics of this protein, but was unable to delve into its folding dynamics and hence to fully characterize its properties and possible implications for bacterial adhesion. Here, with magnetic tweezers, we have fully addressed the dynamics of FimA. Around 100-150 FimA subunits compose the structure of a mature type 2 pilus^60,61^. The isopeptide bonds in FimA prevent the mechanical unfolding of most of the protein, and only the IDL motif extends at very high forces. Under mechanical stress, the multimodular structure of the type 2 pilus would extend as the IDLs are unfolded, reducing the tension at the pilus tip-adhesin/ligand interaction and preventing the mechanical rupture of the pilus backbone. From the chemical point of view, the oral cavity offers an aggressive cocktail of antimicrobial compounds that can target and weaken pilus stability while the IDLs remain extended, such as proteases and peroxidases^62^. From the mechanical perspective, successive shocks would end up cleaving the pilus unless IDL contraction occurs and mechanical stability is restored^63^. Hence, folding on short timescales once the mechanical stress ceases—like in between mastication cycles—is mandatory to prevent pilus chemical and mechanical damage. The fact that the force ranges where FimA folding and unfolding occur are so separated enables the dissipation of massive amounts of heat. Because of the multimodular structure of the type 2 pili, the folding-unfolding of its hundreds of FimA subunits would make each pilus to operate as a shock absorber that damps the mechanical stresses that challenge bacterial adhesion.

The concept of energy dissipation through force-induced bond rupture and molecular extension has been previously proposed for tenascin^64^, bone^65,66^, and spider silk^67^. In these works, the structures under study were exposed to cycles of extension and retraction to trigger bond rupture and reformation, respectively. In all cases, molecular extension required high forces, while retraction and native structure reestablishment occurred at very low force, which generated a large hysteresis that, hence, indicated the large amounts of energy dissipated during this process. These sacrificial bonds, which are responsible of the mechanical stability of these structures, break at high force and are restored once the mechanical stress is removed and entropic forces collapse the extended structure. In *A. oris* type 2 pilus, these sacrificial bonds would be the interactions that prevent IDL extension and hold FimA native structure, which putatively could be placed in the tightly closed interdomain surface that exists between the N2 and N3 domain of FimA^46^. This interdomain structural arrangement is a signature feature of Gram-positive pilins, which could be indicative of a widespread biomechanical strategy in this bacterial group. The role of proteins as mechanical buffers or shock-absorbers has been suggested not only for bacterial adhesion structures^68,69^ or tenascin^64^, but also for proteins such as utrophin^70^, dystrophin^71^, or α-actinin 1^72^, which indicates that this nanomechanical property could be widespread among proteins with mechanical roles.

To reach larger forces in magnetic tweezers while maintaining unaltered its resolution constitutes a technical feat, which will allow the study with unprecedented precision of biomolecules and structures characterized by their mechanical sturdiness and potential dissipative capacities. The fact that it is also possible to study with high precision the dynamics of the mechanosensitive R3 talin domain, without any resolution loss, indicates that subtle force changes can be applied, which highlights the M-450 probes as the perfect tool for investigating a vast number of molecular processes that occur along a large spectrum of force magnitudes.

## Supporting information

Suppementary Information

## Acknowledgements

This research was supported by the National Institutes of Health grant R35129962 (J.M.F). A.A-C. expresses his gratitude to Fundación Ramón Areces (Madrid, Spain) for financial support. Correspondence and request of material should be addressed to A.A-C..

## Author contributions

A.A-C. and J.M.F designed the research. C.L.B and A.A-C designed and developed the split-protein contructs and the solid-phase procedure. A.A-C carried out the experiments. A.A-C and R.T-R. analyzed the data. A.A-C. and J.M.F. wrote the manuscript.

