## Supplementary material for "Magnetic tweezers meets AFM: ultra-stable protein dynamics across the force spectrum": Suppementary Information

### SUPPLEMENTARY INFORMATION

**Table of contents**

**Page 2 – Supplementary Tables**

**Pages 3-8 – Supplementary Figures**

**Pages 9-14 – Supplementary Methods**

**Page 15 - References**

**Supplementary Table 1**

The table below contains the values of the parameters obtained from the simulateneous fit to Eq. 2 (FJC) and Eq. 3 (Bell). The Global Fit procedure from Igor Pro 8.04 (Wavemetrics) was employed.

| Parameter | Value |
|---|---|
| $\Delta L_c$ (Eq. 2, protein L) | 14.67 ± 0.82 nm |
| $l_k$ (Eq. 2, protein L) | 1.23 ± 0.50 nm |
| $\Delta L_c$ (Eq. 2, I27) | 23.17 ± 0.98 nm |
| $l_k$ (Eq. 2, I27) | 1.50 ± 0.63 nm |
| $\ln k_U^0$ (Eq. 3, I27) | -7.24 (fixed value from reference[1]) |
| $x_U^{\dagger}$ (Eq. 3, I27) | 0.19 nm (fixed value from reference[1]) |

**Supplementary Table 2**

The table below contains the values of the parameters obtained from the simultaneous fit to Eq. 5 (FJC) and Eq. 6 (Bell). The Global Fit procedure from Igor Pro 8.04 (Wavemetrics) was employed.

| Parameter | Value |
|---|---|
| $\Delta L_c$ (Eq. 5, protein L) | 14.07 ± 0.53 nm |
| $l_k$ (Eq. 5, protein L) | 1.11 ± 0.16 nm |
| $\Delta L_c$ (Eq. 5, I27) | 23.79 ± 0.38 nm |
| $l_k$ (Eq. 5, I27) | 0.95 ± 0.15 nm |
| $\ln k_U^0$ (Eq. 6, I27) | -7.24 (fixed value from reference[1]) |
| $x_U^{\dagger}$ (Eq. 6, I27) | 0.19 nm (fixed value from reference[1]) |

#### Supplementary Figures

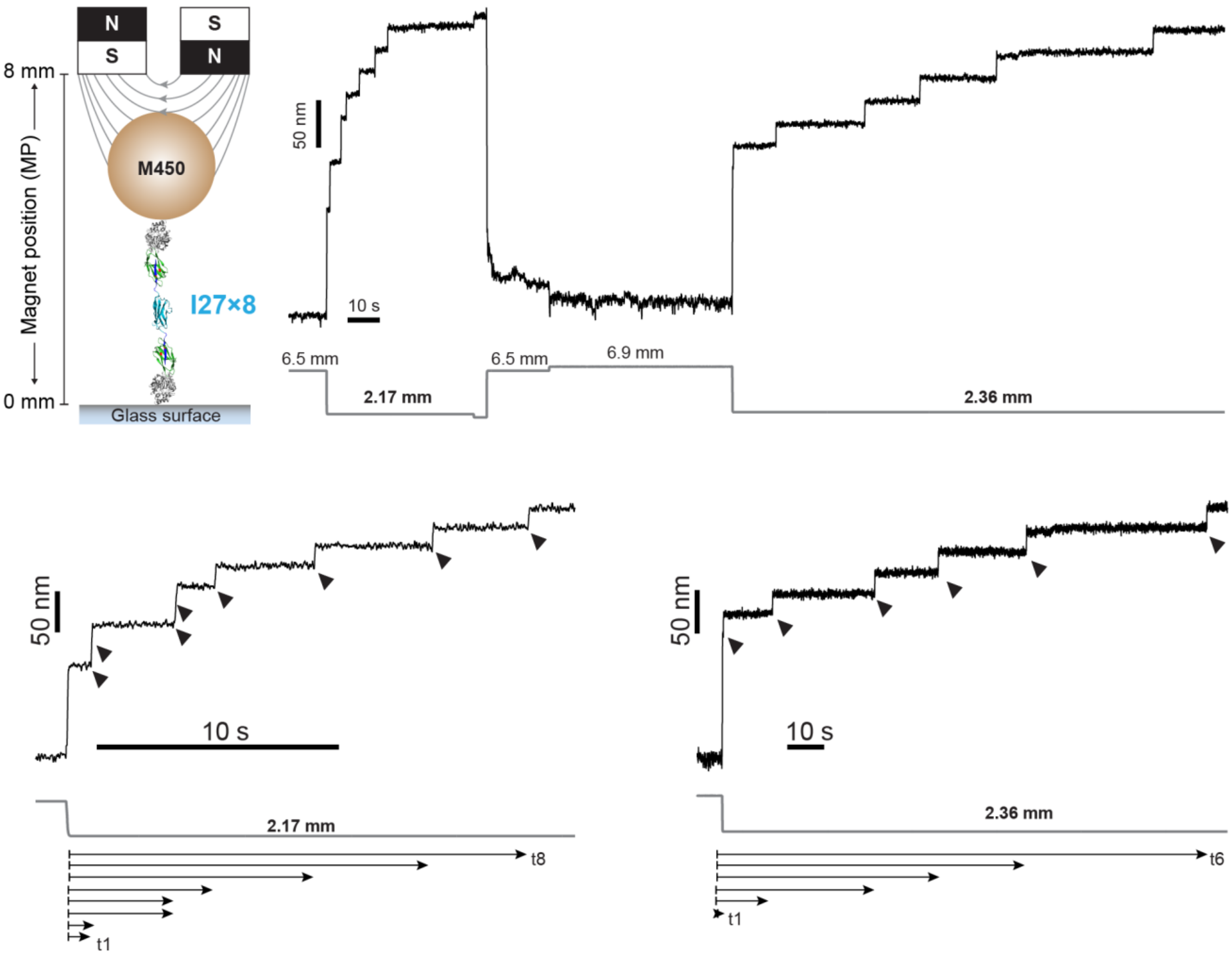


**Supplementary Figure 1. Unfolding dwell-times of I27 in the permanent magnets configuration.** Trajectory of an I27 octamer in the permanent magnets configuration. The unfolding kinetics of I27 are calculated at different magnets positions; here, at 2.17 mm and 2.36 mm, which in the magnet law correspond with 114 pN and 99 pN, respectively. Below are detailed the trajectories with the I27 unfoldings (8 domains at 2.17 mm, and 6 domains at 2.36 mm) and how their dwell-times (t1, t2, etc.) were collected.

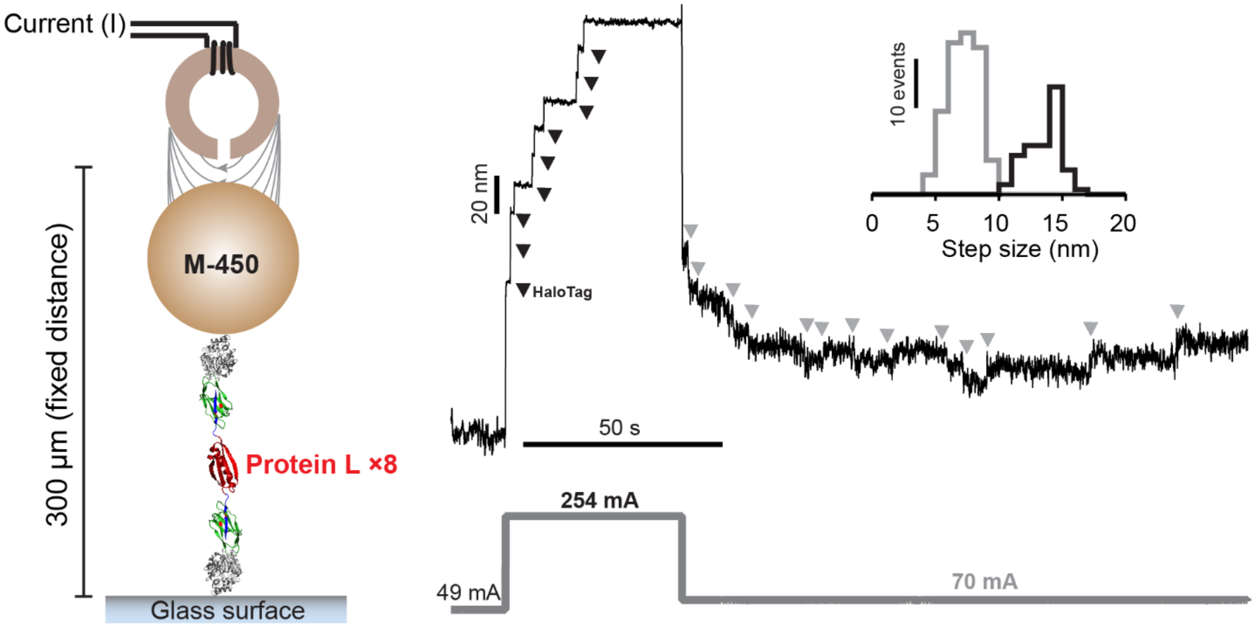


**Supplementary Figure 2. Protein L octamer force-dependent (un)folding step sizes in the electromagnetic tape-head configuration. a)** Force-clamp trajectory of an octamer of protein L. Along the experiment, the current is modified, which triggers protein unfolding (current value from 49 mA to 254 mA, black arrows), which results in step sizes of 13.5±1.4 nm (mean±SD. Inset, black histogram, n=58), and folding transitions (current value from 254 mA to 70 mA) which yield 7.1±1.2 nm step sizes (inset, grey histogram, n=122).

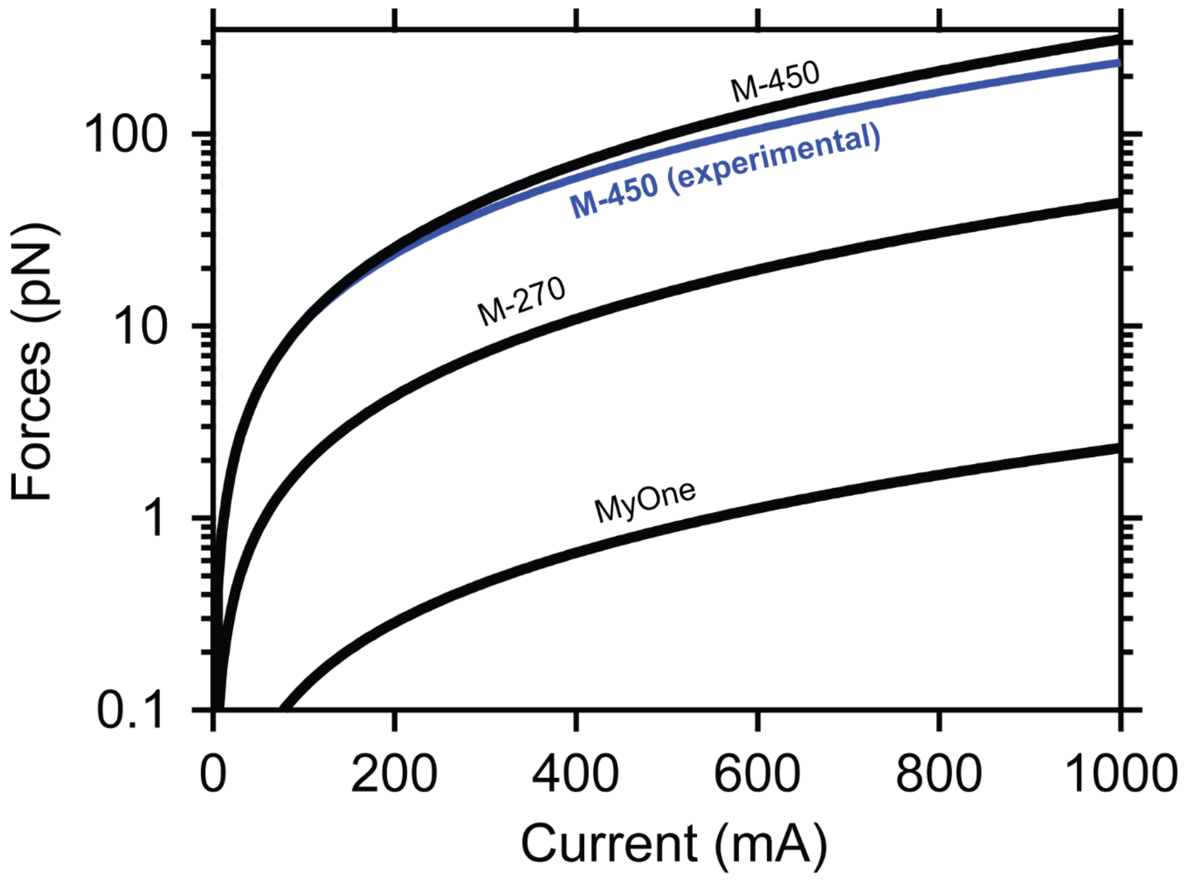


**Supplementary Figure 3. Theoretical current law for different superparamagnetic beads.** Force range prediction (black lines) as a function of the current for MyOne (*d* = 1 µm), M-270 (*d* = 2.8 µm), and M-450 (*d* = 4.5 µm) beads. Blue line represents the current law determined experimentally with the M-450 beads. **Supplementary Methods** contain the calculations for determinining the *A* and *B* parameters of the current laws specific for each of the different beads.

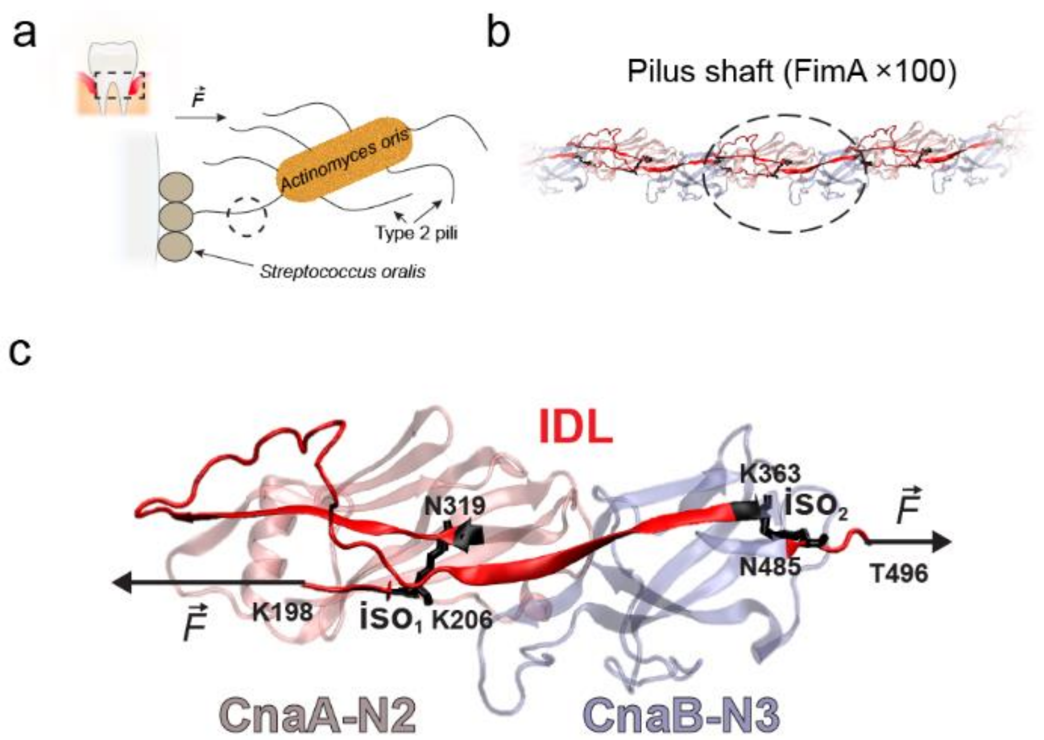


**Supplementary Figure 4. Gram-positive oral pathogen *Actinomyces oris* utilizes the type 2 pili to establish interspecies interactions. a**) Pathogenic bacteria colonize the teeth surface and cause dental plaque. Despite mechanical challenges such as mastication or teeth brushing, *Actinomyces oris* is able to attach and establish cell-to-cell interactions with other pathogens (*Streptococcus oralis*) using its type 2 pili, the micrometer-long structures used for adhesion. **b)** *A. oris* type 2 pilus shaft is formed by the covalent concatenation of hundreds of FimA pilin proteins. In the pilus, each FimA subunit is connected to the next one through intermolecular isopeptide bonds formed between T496 residue (domain CnaB-N3) of the previous FimA and the K198 residue (CnaA-N2 domain) of the next one (see panel **c**). **c)** FimA is composed by 3 Ig-like domains—from N to C-terminus: CnaA-N1 (not shown), CnaA-N2, and CnaB-N3. Each of these domains contain intramolecular isopeptide bonds, and the polypeptide sequence trapped between the CnaA-N2 ($iso_1$, K206-N319) and CnaB-N3 ($iso_2$, K363-N485) isopeptide bonds defines a structural motif termed isopeptide-delimited loop (IDL, highlighted in red). Due to the location of the isopeptide bonds and the head-to-tail linking between FimA subunits, the force is propagated along the pilus through the interconnected IDL motifs, which are the only structure of the protein and from the pilus backbone that can extend under force (PDB: 3QDH).

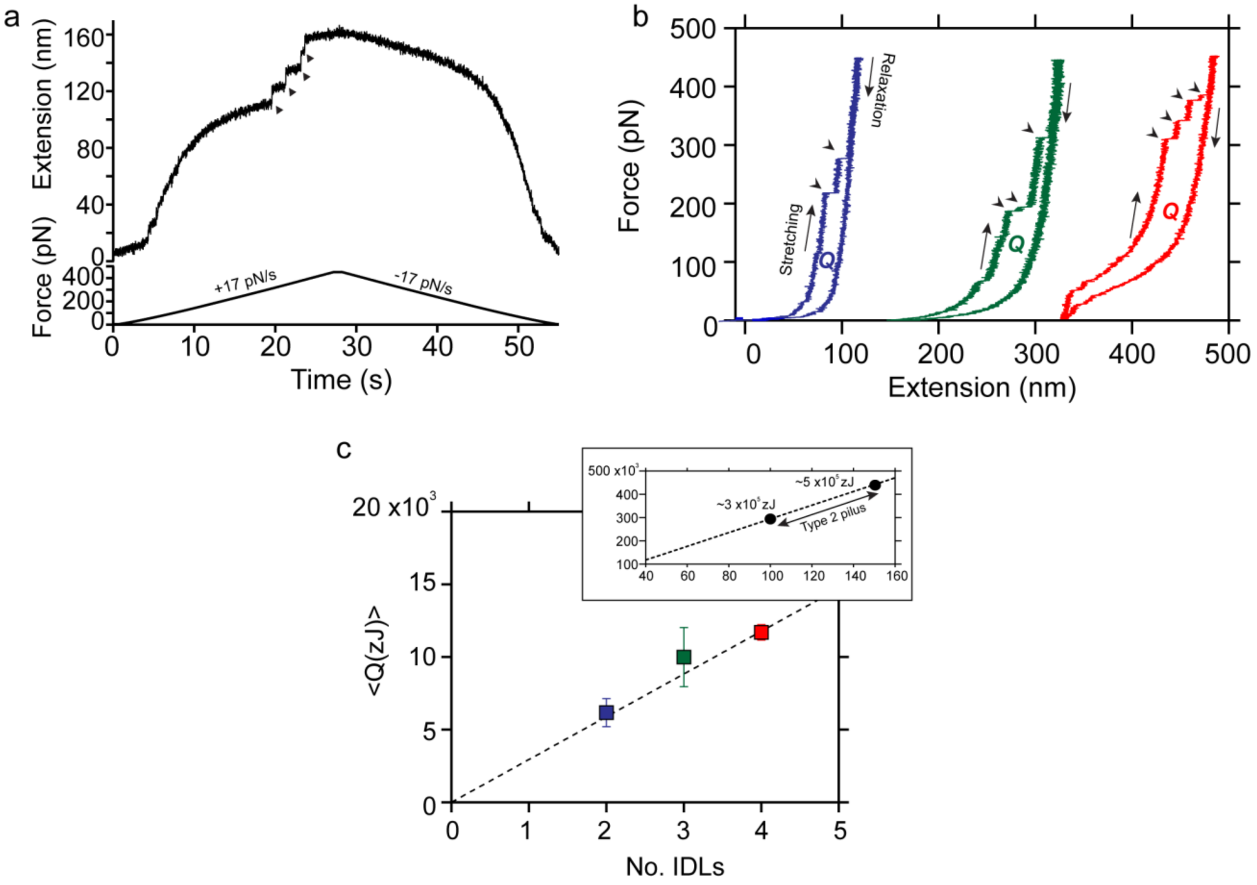


**Supplementary Figure 5. Energy dissipation of FimA. a)** Force-ramp experiment on a polyprotein composed of four FimA subunits. Force is increased from 4 pN to 450 pN at a constant rate of 17 pN·s-1. Once the maximum force is reached, the force is decreased to 4 pN at the same loading rate. Black arrows indicate the unfolding of the 4 IDLs present in the polyprotein. **b)** Force vs Extension plots of stretching-relaxation cycles (±17 pN·s-1) in polyproteins exhibiting the unfolding of 2 (blue), 3 (green), and 4 (red) IDLs. The area enclosed by the stretching and relaxation curves is the amount of heat (*Q*) dissipated in these trajectories. **c)** Average heat dissipated (<*Q*>) as a function of the number of unfolding IDLs (2 IDLs, n=8; 3 IDLs, n=4; 4 IDLs, n=3). The dotted line represents a fit of the data using a model (see **Supplementary Methods**) that predicts a linear relationship between the average heat dissipated and the number of IDLs. Inset shows an extrapolation to the heat values that a 100-150 FimA pilus could dissipate.

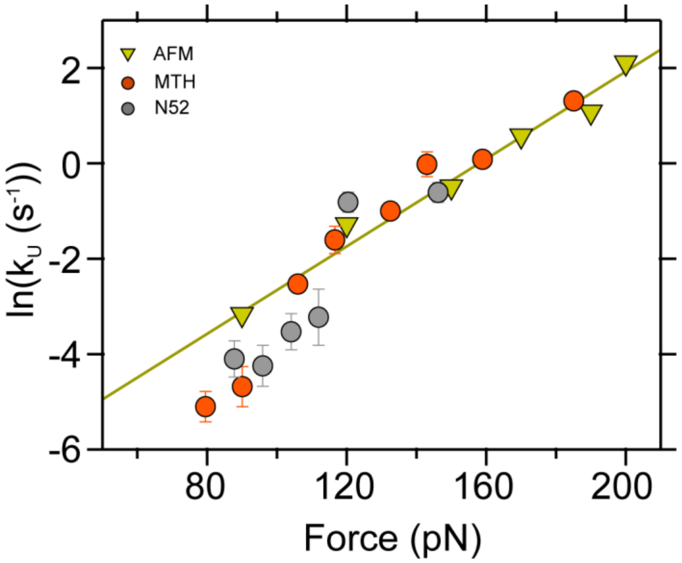


**Supplementary Figure 6. Comparison of the I27 unfolding kinetics under force with AFM and magnetic tweeezers.** I27 unfolding kinetics in AFM (yellow arrows) and in the two magnetic tweezers configurations (MTH: magnetic tape head, orange circles; N52: permanent magnets, grey circles) employing the superparamagnetic beads M-450. The line represents a Bell model fit to the AFM measurements from reference 1 ($\ln k_U{}^0$=17.24, $x^\dagger$=0.19 nm).

#### Supplementary Methods

##### Protein engineering and expression

All the reagents employed in this research were from Sigma-Aldrich, unless otherwise specified. The polyproteins SnoopTag-(protein L)$_8$-SpyTag, SpyTag-(I27)$_8$-SpyTag, and SpyTag-(FimA)$_4$-SpyTag were cloned into the pQE80L expression plasmid, the anchor proteins SnoopCatcher-HaloTag, SpyCatcher-HaloTag, and HaloTag-Talin R3$^{IVVI}$-(Spy0128)$_2$-SpyTag were cloned into and expressed from a modified pFN18a-HaloTag®T7 Flexi® vector (provided by R.T. Sauer from Massachusetts Institute of Technology). All of the protein constructs contain a 6xHistag for the purification procedure. All the cloning steps were done following the strategy previously described[2]. In brief, all the genes encoding the proteins (except HaloTag) contain a 5' *BamHI* and a 3' *BglII* restriction sites. Upon digestion with these two enzymes and purification of the genes, these are ligated into the pFN18A or the pQE-80L plasmids which have been previously digested with BamHI. In pFN18A the *BamHI* restriction site is placed on the 3' of the sequence of HaloTag and the TEV site, while in the modified pQE-80L is placed between the SpyTag peptide sequences (or in between the SnoopTag and SpyTag sequences). BamHI and BglII produce compatible ends and after ligation, the gene can be inserted in the correct or incorrect orientation in the plasmid. The correct insertion of the gene is identified with digestion with BamHI and KpnI (pQE-80L) or BamHI and BlpI (pFN18A), and sequencing. BlpI restriction site is located in the plasmid sequence after the stop codon. In the correct orientation, this digestion releases two DNA fragments corresponding with the empty plasmid and the gene. If the insertion was in the incorrect orientation, the digestion produces one fragment of a few pair of bases (<50 bp) and a large DNA fragment corresponding with the sequence of the plasmid plus the gene. All the cloning and amplification steps were conducted on the *Escherichia coli* XL10-Gold strain (Agilent Technologies).

*E. coli* ERL strain cells (R.T. Sauer) were transformed with the plasmids containing the chimeric proteins. Protein expression induction and purification protocols were done as previously described[3]. In brief, the transformed cells were grown at 37°C with 250 rpm constant shaking until $OD_{600}$>0.6. Expression was induced with 1 mM IPTG at 25°C with 250 rpm constant shaking for 12-16 h. Protein extraction from cells was achieved through mechanical lysis with a French press (Sim-Aminco), and the protein isolation from the lysate was done with the His60 Ni Superflow

Resin (Clontech). Further purity was achieved with a size exclusion chromatography step (Superdex 200 FPLC column, GE Healthcare). Proteins were eluted in 10 mM Hepes (pH 7.2), 150 mM NaCl, 1 mM EDTA and stored with 10% v/v of glycerol at -80°C until use.

#### Glass and bead surfaces functionalization

Tosyl-activated Dynabeads® M-450 (Thermo Fisher Scientific) were functionalized with a 3 µM:1 µM mixture of the HaloTag protein and HaloTag-SpyCatcher, respectively. Beads were incubated at 4°C and at 18 rpm constant rotation (Labnet), and maintained under these conditions until their use. Before experiments, excess unbound proteins were removed by iterative pelleting of the M-450 beads, supernatant removal, and resuspension in PBS buffer. Fluid chamber assembly and glass surface functionalization were done as previously described[4].

#### Double covalent and molecular assembly

The fluid chambers were incubated with a 3-5 nM concentration of the anchor protein HaloTag-SpyCatcher, or HaloTag-SnoopCatcher, at room temperature for 30 min. In the case of the protein of interest talin $R3^{IVVI}$ domain, the construct HaloTag-$R3^{IVVI}$-$(Spy0128)_2$-SpyTag was directly coupled to the surface. After, the fluid chambers were rinsed extensively with PBS to remove not bound protein, and immediately after the protein of interest flanked by the peptides SnoopTag-SpyTag or SpyTag-SpyTag was incubated at 5-10 µM at room temperature for 1 h. Fluid chambers were extensively rinsed again with PBS buffer supplemented with 10% of ascorbic acid (pH 7.4). The fluid chamber containing the first two layers of the molecular assembly was then placed on the microscope stage (permanent magnets) or at the base of the card-reader piece (electromagnetic tape-head). Once there, the SpyCatcher-HaloTag-functionalized M-450 beads were added to the fluid chamber and incubated for 3 minutes, to close the assembly. Immediately after, the magnets were approached (permanent magnets) or the current was turned on (electromagnetic tape-head) to exert an initial force of 4 pN and start the experiments.

#### Magnetic tweezers configurations

Experiments were conducted on custom-built magnetic tweezers apparatuses, as previously described[4,5].

The permanent magnets configuration chassis consists of an inverted microscope (Olympus IX-71/Zeiss Axiovert S100). Reference and magnetic beads are illuminated with a collimated cold white LED source (ThorLabs), and visualized with a 63X or 100X oil-immersion plan apochromat objective (Zeiss/Olympus). The objective position, and hence the focal plane, is controlled with a P-725 nanofocusing piezo actuator (Physik Instrumente). A CMOS Ximea MQ013MG-ON camera was used for image acquisition, and the image processing was realized with a custom program written in C++/Qt. The NI USB-6289 multifunction DAQ card (National Instruments) was used for controlling the piezo position and the magnets distance and acquire the data. Magnets position were controlled with a voice coil actuator (LFA-2010, Equipment Solutions) operate under electronic feedback with a custom-made PID controller.

The magnetic tape head configuration operates with the same components as the permanent magnets one, but instead of having a microscope chassis it has all of the components arranged vertically along the scaffold provided by a 95 mm construction rail (500 mm height, X95-500, ThorLabs). The optical path and components are held along the vertical axis of one side of the tower with clamps for optical rail carriage (X95 Series, Newport). The electromagnetic tape head (Brush Industries, 902836) is held fixed on a custom-designed CNC aluminum piece (card-reader component in **Fig. 4a**) designed in Autodesk Fusion 360 (Adobe). The base of the card-reader, which is where the fluid chamber lies, has an opening that permits to accommodate the objective to visualize the reference and magnetic beads. The 25 µm gap of the tape head is aligned to the center of the optical path and its vertical distance from the upper surface of the bottom glass is 300 µm. The fluid chamber is held with a custom-made metal stamped fork whose lateral arms are coated with double-sided tape (Scotch). The unit formed by the fork and the fluid chamber is moved along the X and Y directions with a XY Linear Stage (NewPort). A thorough description of this setup and all the elements that compose it will be reported in a paper that is currently under preparation.

#### Analysis

The analysis was done with Igor Pro 8.0 software (Wavemetrics). Recordings were smoothed using a 4th order Savitzky-Golay filter with a box size of 101 points. Step sizes were determined by measuring the distance between the peaks of Gaussian fits done on the unfolding steps. FimA folding probability was calculated as the ratio between the number of unfolded domains and the number of domains able to fold after 100 s at each of the forces tested. A jackknife estimator was used for the calculation of the average probability and the standard deviation. Talin $R^{IVVI}$ domain folding probability was calculated as the ratio of time spent in the folded and unfolded states as a function of the force.

#### Theoretical current law for different types of beads in the magnetic tape head configuration

As described in the main text, the fixed distance between the magnetic tape head and the bead ($z$ = 300 µm) simplifies the calculation of the law that describes the force as a function of the current (Eq. 4, main text). This situation leaves the force law depending on the current value and the parameters $A$ (Eq. 1) and $B$ (Eq. 2). As we described previously[5], the value of these parameters is specific of the magnetic tape head and the superparamagnetic beads employed during the calibration.

$$A = 8V\chi_b\mu_0 \frac{N^2}{g^3}\frac{\eta^2}{\pi^2} \qquad \text{Eq. 1} \qquad\qquad B = 4\rho V\mu_0 \frac{N}{g^2}\frac{\eta}{\pi}M_{0x} \qquad \text{Eq. 2}$$

In $A$ and $B$, the bead-specific parameters are the volume ($V$), the initial susceptibility of the bead ($\chi_b$), the density ($\rho$), and the initial magnetization ($M_{0x}$). Since we want to calculate a theoretical force-current law for different beads under the same magnetic tape head, we can use the properties and the current law of the M-270 beads[5] to calculate the value of the $A$ and $B$ parameters for other beads such as MyOne or M-450. We use the values of the properties of these beads reported by the thorough characterization of Fonnum et. al (2005)[6] to calculate their corresponding force-current law. As an example, below we show the calculations for the M-450 beads (shown in **Supplementary Figure 3**). The theoretical value of $A$ for the M-450 beads ($A'$) can be obtained as:

$$\frac{A}{A'} = \frac{V}{V'}\frac{\chi_b}{\chi'_b} \qquad \text{Eq. 3} \qquad\qquad A' = \frac{V'}{V}\frac{\chi'_b}{\chi_b}A \qquad \text{Eq. 4}$$

Where $A$ (2.8×10$^{-5}$ pN/mA$^2$), $V$ (1.1×10$^{-17}$ m$^3$), and $\chi_b$ (0.8) are the values from the M-270, and $A'$, $V'$ (4.8×10$^{-17}$ m$^3$), and $\chi_b'$ (1.6) are the values for the M-450. The value of $\chi_b$ and $\chi_b'$ is dimensionless and it is the product of the bead density ($\rho$, kg·m$^{-3}$) and the initial magnetic susceptibility ($\chi$, m$^3$·kg$^{-1}$)

$$\chi_b = \chi\rho \qquad \text{Eq. 5}$$

Therefore, the theoretical value of $A'$ for the M-450 beads is 2.6×10$^{-4}$ pN·mA$^2$. In the case of the $B$ parameter, we follow the same procedure:

$$B' = \frac{\rho'}{\rho}\frac{V'}{V}\frac{M'_{0x}}{M_{0x}} \qquad \text{Eq. 6}$$

From where we obtain $B'$=7.9×10$^{-2}$ pN·mA$^{-1}$.

The theoretical values of both parameters $A$ and $B$ are very close to the experimental ones, which indicates the strength of our calibration method based on protein extension changes and unfolding kinetics under force. Moreover, it corroborates our magnetic tape head tweezers as a solid framework for the calibration of other superparamagnetic probes.

**Derivation of the average heat dissipated by an unfolding polyprotein**

We assume that the FimA polyprotein chain has $N$ domains each with a folded extension of $\Delta$ and an unfolded contour length of $L_c$. Also, we describe its extension-force relation using the Freely-Jointed Chain model[7]. Hence, the extension of the folded protein $x_F$ and of the unfolded protein $x_U$ can be written as:

$$x_F = L_F\left[\coth\left(\frac{Fl_k}{k_BT}\right) - \frac{k_BT}{Fl_K}\right] \qquad (1)$$

$$x_U = L_U\left[\coth\left(\frac{Fl_k}{k_BT}\right) - \frac{k_BT}{Fl_K}\right] \qquad (2)$$

where $l_K$ is the Kuhn length of the folded and unfolded polyproteins, which we assume to be equal, and $L_F$ and $L_U$ the contour length of the folded and unfolded chain, this is:

$$L_F = L_0 + N\Delta \quad (3)$$

$$L_U = L_0 + NL_C \quad (4)$$

being $L_0$ the contribution of the contour length of unstructured linkers.

The polyprotein experiences some mechanical perturbation, such as a fast force ramp or force shock that unfolds its $N$ domains. We want to calculate the average heat dissipated $\langle Q \rangle$ in this process. To simplify, we assume that this average heat is equal to having all domains unfolding at an average $F_U$, which is a the average unfolding force of the FimA domains when pulled at a certain rate in force-ramp. Then, the average dissipated heat can be calculated as:

$$\langle Q \rangle = \int_0^{F_U} [x_U(F) - x_F(F)]dF \quad (5)$$

where we assume that the domains refold at 0 force or at a very low force ($F_U >> 0$) when the mechanical shock is relaxed. This integral can be solved analytically, and obtain an analytical expression for the average dissipated heat:

$$\langle Q \rangle (F_U, N) = N \frac{\Delta L_c k_B T}{l_K} ln \left[ \frac{k_B T sinh\left(\frac{F_U l_K}{k_B T}\right)}{l_K F_U} \right] \quad (6)$$

where $\Delta L_c = L_c - \Delta$.

A few observations on this expression. First, the dissipated heat scales linearly with the number of domains of the polyprotein. We are assuming that all domains unfold during the mechanical shock. In case that is not true, we would have ($N_U L_c$ - $N\Delta$), still a linear scaling. Second, the initial extension does not affect to the dissipated heat; it extends and relaxes in equilibrium. Third, this value can be calculated for a given experiment. $\Delta L_c$ and $l_K$ are obtained from measurement of the step-sizes; we can assume $l_K$ to be the same for the folded and unfolded polyprotein since soft linkers dominate. $F_U$ can be measured as the average unfolding force for a given condition, for instance for a certain pulling rate. In case we know the geometric parameters of the protein ($k_U^0$ and $x_U^{\dagger}$), $F_U$ can be calculated for arbitrary experimental conditions from a Bell-like expression.
